# CryoDiff: An uncertainty-aware diffusion model for Cryo-EM map enhancement

**DOI:** 10.64898/2026.06.04.730282

**Authors:** Bocheng Wen, Bintao He, Yiran Cheng, Shaozhen Zhou, Fa Zhang, Renmin Han

## Abstract

Cryogenic electron microscopy (cryo-EM) enables high-resolution structural determination of large macromolecular complexes. However, the interpretability of cryo-EM maps is often hindered by substantial background noise and signal attenuation, which obscure structural details. Although existing post-processing methods can partially mitigate these artifacts, they typically suffer from over-smoothing and lack reliable confidence estimation. Here, we present CryoDiff, an uncertainty-aware diffusion model for cryo-EM map enhancement. CryoDiff employs a multi-step diffusion process to progressively denoise and restore high-resolution structural features. Importantly, CryoDiff incorporates a voxel-wise confidence metric derived from Monte Carlo sampling. It unifies map enhancement and voxel-level uncertainty estimation within a diffusion-based generative framework, representing the first approach to achieve such joint modeling for cryo-EM map enhancement. In comprehensive experiments, CryoDiff markedly out-performs existing methods in both map–model correlation and map interpretability, improving the average *FSC*_0.5_ metric by 0.356 Å over state-of-the-art approaches. When applied to *de novo* model building with ModelAngelo, CryoDiff further increases model completeness by 5.5%, exceeding the gains achieved by competing method.

## 1 Introduction

Cryogenic electron microscopy (cryo-EM) has emerged as a powerful technique for determining the structures of macromolecular complexes in their native states. In the single-particle cryo-EM workflow, three-dimensional density maps are reconstructed from tens of thousands of particle images (Nogales and Scheres, 2015) to serve as a reference for atomic model building. However, raw experimental maps frequently suffer from contrast loss and high-frequency signal attenuation (Rosenthal and Henderson, 2003) due to factors such as intrinsic structural flexibility, radiation damage (Glaeser and Taylor, 1978), and cumulative errors during the 3D reconstruction process. Such signal degradation obscures critical structural features, introducing significant ambiguity into backbone tracing and side-chain assignment. Consequently, various post-processing approaches have been developed to recover these missing details and maximize map interpretability.

Traditional map sharpening methods aim to compensate for the contrast loss in cryo-EM maps by amplifying high-frequency Fourier components, which is typically achieved either by applying a global B-factor (Kimanius *et al*., 2016; Zivanov *et al*., 2018; Terwilliger *et al*., 2018) or employing local sharpening techniques that leverage prior information (Ramírez-Aportela *et al*., 2020; Jakobi *et al*., 2017). In addition, density modification methods improve cryo-EM maps by incorporating prior knowledge, such as solvent characteristics or local noise distributions (Terwilliger *et al*., 2020a,b). These approaches employ physically or statistically grounded algorithms to enhance cryo-EM maps, providing consistent and reliable processed densities. Consequently, these methods have been widely adopted by the cryo-EM community as standard procedures in the post-processing of cryo-EM maps, serving as important steps for both map deposition and downstream structural analysis. Despite their widespread adoption, these conventional approaches suffer from notable limitations. Global B-factor-based sharpening cannot account for spatially varying signal-to-noise ratios, often leading to over-sharpening in flexible regions and under-sharpening in stable cores. Meanwhile, the efficacy of many local sharpening methods is restricted by their reliance on additional inputs, such as segmentation masks or pre-existing atomic models, limiting their applicability to novel structures where such priors are unavailable or unreliable.

Deep-learning-based methods have been proposed for automatic cryo-EM map enhancement. Early attempts like DeepEMhancer (Sanchez-Garcia *et al*., 2021) employs 3D U-Nets to replicate enhanced map sharpened by traditional model LocScale (Jakobi *et al*., 2017). To overcome such dependency, some works (Subramaniya *et al*., 2021; He *et al*., 2023; Cao *et al*., 2025; Zhang *et al*., 2025; Selvaraj *et al*., 2024) were trained on paired data consisting of noisy experimental maps and noiseless maps simulated from high-quality atomic models. Among these approaches, EMReady2 (Cao *et al*., 2025) stands out as the leading post-process tool, which using heterogeneity-aware deep learning to improve Cryo-EM maps. However, EMReady and similar deterministic deep-learning-based approaches exhibit critical limitations. First, these deep-learning approaches operate as “black boxes” that directly modify real-space densities without providing explicit explanations or quantitative confidence measures for the enhanced features. Second, cryo-EM reconstructions inherently represent an ensemble of conformations due to molecular flexibility (Herzik *et al*., 2019). By relying on single-pass feedforward networks that regress to a single deterministic output, these methods tend to produce overly smoothed densities, often blurring high-frequency details.

Here we propose CryoDiff, an uncertainty-aware diffusion model for cryo-EM map enhancement that employs a conditional denoising diffusion probabilistic model (Ho *et al*., 2020). Instead of regressing to a single solution, CryoDiff learns the distribution of high-quality structures and iteratively refines details from random Gaussian noise. This gradual denoising framework provides two crucial advantages over deterministic methods. First, it naturally mitigates over-smoothing by avoiding direct regression to a single, averaged solution. Second, by modeling the full conditional distribution, CryoDiff can generate a diverse ensemble of plausible reconstructions, thereby capturing the inherent uncertainty of the enhancement process. To quantify this uncertainty, CryoDiff further introduces a novel voxel-wise reliability metric, which measures confidence in the recovered features by performing Monte Carlo sampling and evaluating density variations across the ensemble.

To evaluate the performance of CryoDiff, we conducted comprehensive experiments on a large-scale benchmark comprising 143 primary maps and 80 pairs of half maps with resolution ranging from 2 to 5 Å. The case results show that CryoDiff produces maps with substantially enhanced interpretability, characterized by improved structural continuity and the recovery of intricate side-chain densities often obscured in experimental maps. We quantitatively assessed its performance using map-model correlation metrics, including map-model *FSC*_0.5_, Q-score(P), Q-score(N), *CC*_box_, *CC*_mask_ and *CC*_peaks_. Specifically, CryoDiff boosts the average *FSC*_0.5_ resolution to 3.045 Å (primary) and 2.946 Å (half-maps), outperforming the state-of-the-art method by 0.356 Å and 0.337 Å, respectively. Furthermore, this superior map quality enables downstream tools, like phenix.map to model and ModelAngelo, to generate more accurate and complete *de novo* atomic models. Notably, the atomic model built by ModelAngelo from the CryoDiff-processed maps achieved an average completeness of 79.3%, representing an improvement of 5.5% compared with the deposited maps.

## 2 Results

### 2.1 Overview of CryoDiff

The complete workflow of CryoDiff, illustrated in Figure 1, consists of four main stages: data generation, model training, reverse inference, and uncertainty estimation. In the data generation stage, we first collected 980 experimental cryo-EM maps from the Electron Microscopy Data Bank (EMDB) and split them into training (737), validation (100), and test (143) sets. Subsequently, for the training set, simulated maps were generated from corresponding PDB (Berman *et al*., 2000) models to serve as the ground-truth for model training. All maps were finally uniformly processed by resampling to the voxel size of 1 Å, normalizing intensity values to the range [0, 1], and partitioning into boxes of size 48^3^. In the model training stage, CryoDiff, a diffusion-based 3D U-Net (detailed in 3.3 and Supplementary Figure S4) (Rombach *et al*., 2022a), is trained to predict the Gaussian noise added to simulated map boxes *x*_0_ at the random diffusion timestep *t*, with the corresponding conditional experimental map box *x*_*c*_, as detailed in Section 3.4. This model is optimized via a hybrid Smooth L1 loss and SSIM loss. In the reverse inference stage, the input map is first partitioned into overlapping boxes of size 48^3^ with a stride of 16. The denoising diffusion implicit model (DDIM) sampler then iteratively generates the enhanced boxes 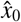 over *S* steps through a reverse diffusion process, which starts from random Gaussian noise and is conditioned on the corresponding input experimental boxes.

**Figure 1:**
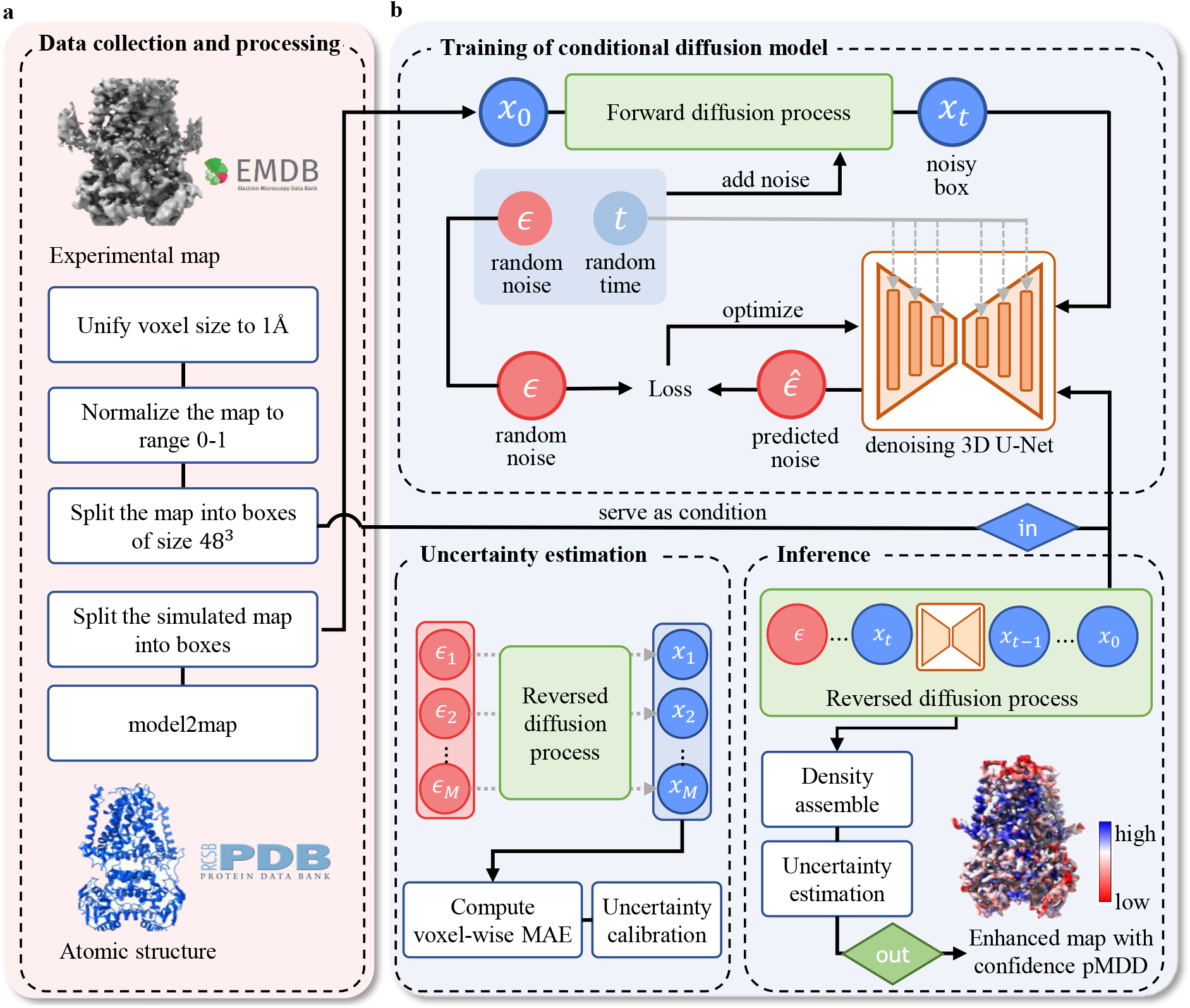
Data processing of maps in CryoDiff. **a** Data preprocessing workflow for training and inference data. **b** Overview of CryoDiff. Supervised training of the conditional diffusion model, CryoDiff, is illustrated in the upper section. For each iteration of training, random Gaussian noise is added to the boxes extracted from the simulated ground-truth map, and the network is trained to denoise them conditioned on the corresponding boxes from the experimental map. The inference of CryoDiff is demonstrated in the lower right section. Reverse diffusion is initiated from random Gaussian noise, conditioned on experimental boxes. The generated boxes are subsequently resampled and assembled into an enhanced map. The lower left section shows the uncertainty estimation process to assess the local quality of the enhanced map. The reverse diffusion process is repeated *M* times to generate an ensemble of predictions. Local uncertainty is quantified as the mean absolute error (MAE) between the ensembles and the enhanced map, which are further calibrated to form a confidence map.

These enhanced boxes are then assembled to form the complete, high-quality enhanced map. In the uncertainty estimation stage, CryoDiff introduces a voxel-wise confidence metric, pMDD, to quantify the structural reliability of the enhanced map, as detailed in Section 3.6.

### 2.2 Local enhanced confidence estimation

Except for the enhancement of cryo-EM densities, the post-processing of cryo-EM reconstructions also involves the validation of the maps. Validation of cryo-EM maps aids the subsequent model-building process by providing quantitative metrics that indicate how likely a feature within the density map represents specific molecular components.

To assess the local reliability of CryoDiff’s enhanced results, we propose a metric termed predicted Multi-Diffusion Difference (pMDD) based on the inherent uncertainty of the diffusion process. The effectiveness of this confidence metric is directly attributed to CryoDiff’s paired training strategy, wherein high-quality map regions correlate with accurate atomic models to provide consistent signals and minimize uncertainty. Conversely, in regions of low quality or structural flexibility, it is inconsistencies between the deposited map and the model-derived map that introduce ambiguity, thus elevating the sampling uncertainty.

The proposed pMDD is initially derived from voxel-wise mean absolute error (MAE) over multiple diffusion samples, leveraging the intrinsic uncertainty of the diffusion process. However, we find that the pMDD–resolvability relationship inferred from raw MAE is influenced by the global resolution of the map, limiting its interpretability across datasets. To address this issue, we introduce an uncertainty calibration strategy that aligns pMDD with established measures of map resolvability. After calibration, pMDD provides a normalized and comparable metric across maps, while preserving the relative ordering of voxel-wise errors. As a result, the calibrated pMDD serves as a reliable proxy for local resolvability and further captures structural variability arising from flexibility, noise, and potential artifacts. Note that, after calibration, higher pMDD values correspond to better local resolvability and higher confidence, aligning its interpretation with Q-score.

To evaluate the reliability of proposed pMDD reliability, we introduced the cryo-EM map of Dnf1 from Saccharomyces cerevisiae (EMD-31487, 3.81 Å) (Xu *et al*., 2022) as a case study (Figure 2). Figure 2a demonstrated the deposited and CryoDiff-processed map colored with pMDD values. In the deposited map, the A and N domains exhibit poor resolution, attributed to their intrinsic flexibility and residual conformational heterogeneity. This underlying instability is also evidenced by their reduced local pMDD values, which are significantly lower than those of the stable transmembrane (TM) domain. Figure 2b illustrates two *α*-helical segments in the TM domain of Dnf1 at different contour level. Unlike EMReady2, which lacks a local quality metric, CryoDiff provides a pMDD-colored confidence map that explicitly highlights regions of structural uncertainty. This crucial feature allows researchers to readily distinguish between well-resolved region and potential artifacts or ambiguous densities. A detailed comparison at the low contour level (black arrows) reveals three key advantages of CryoDiff’s pMDD metric. First, an ASN side chain appears erroneously extended in the EMReady2-processed map, representing a feature that can be easily misinterpreted. In contrast, CryoDiff assigns moderate uncertainty to this region, suggesting that the feature may be misleading while interpretation. Second, the ARG side chain remains ambiguous in both maps, while CryoDiff provides crucial local information by highlighting lower pMDD near the base of the side chain. Third, the pMDD metric effectively identifies residual noise densities which persist in both enhanced maps as highly uncertain regions.

**Figure 2:**
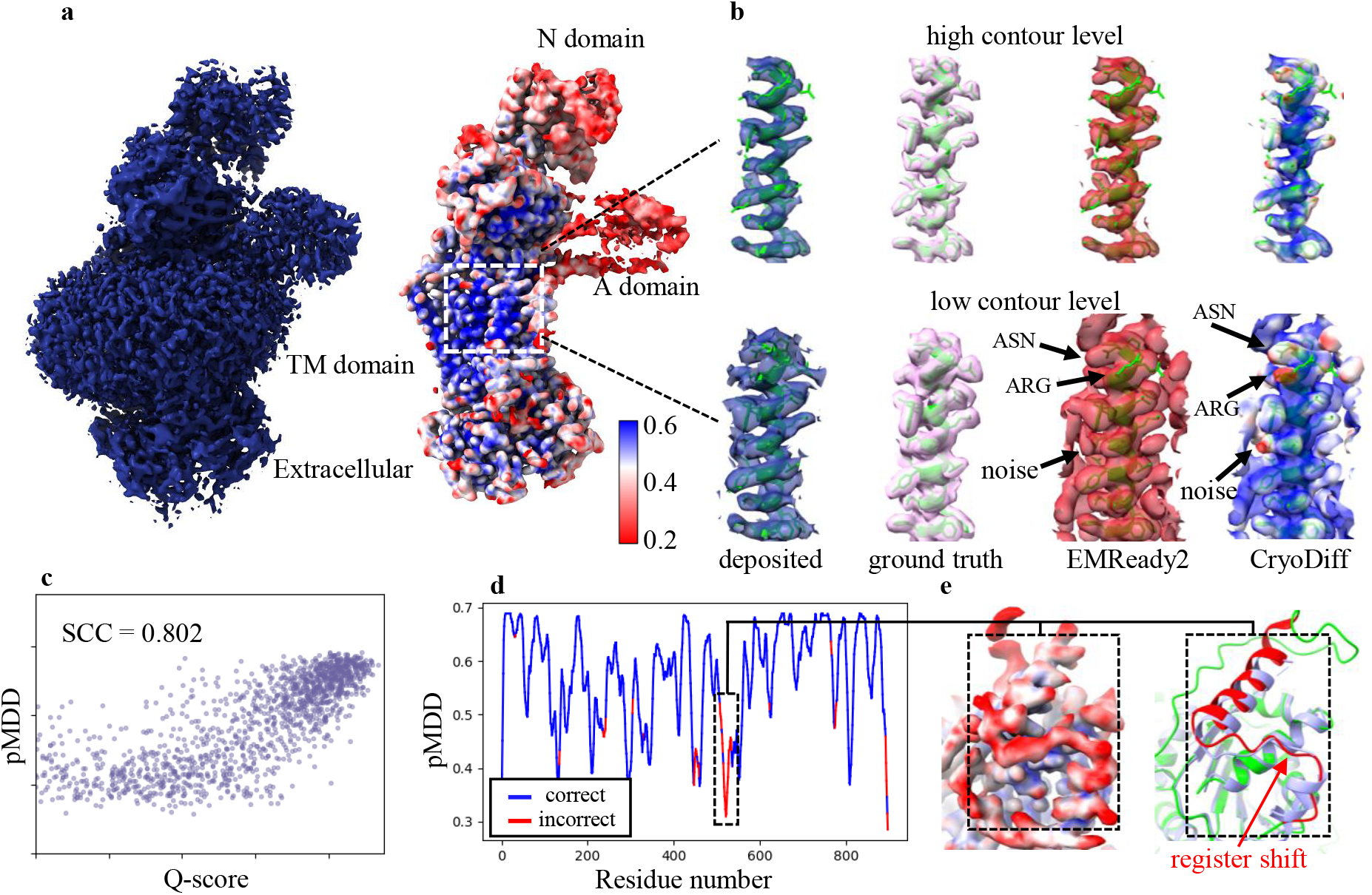
Enhanced result of Dnf1 from Saccharomyces cerevisiae (EMD-31487, 3.81 Å, PDB:7F7F). **a** The deposited map (left), and the CryoDiff-processed map (right) which is colored by pMDD values. **b** Densities of an *α*-helical segment in the TM domain from the deposited map, simulated map, EMReady2-processed map, and CryoDiff-processed map, with contour levels set to enclose an equal volume. Black arrows highlight regions in the CryoDiff map exhibiting high uncertainty. **c** Correlation between residue-wise Q-score and pMDD values. **d** Plot of pMDD over residue number. Residues where the sequence of the deposited model differs from that of the ModelAngelo model are highlighted in red, while residues with matching amino acid types between the two models are shown in blue. **e** Enlarged view of a register-shift region in the deposited PDB model. The CryoDiff-processed map with pMDD is shown on the left. On the right, the deposited PDB structure is displayed with residues involved in the register shift highlighted in red and the remaining residues in green, while the ModelAngelo-predicted model is shown in purple.

To further verify pMDD’s capability to reliably reflect the local resolvability of the enhanced map, we quantified its correlation with the established Q-score values (see Methods 3.9). This analysis yielded a Spearman Correlation Coefficient (SCC) of 0.802 (Figure 2c), confirming that the calibrated pMDD reliably reflects local resolvability measured by Q-score. Moreover, because pMDD reflects local map quality, regions with high pMDD frequently coincide with segments susceptible to modeling errors. As shown in Figure 2d, we plotted pMDD values along the residue index and compared regions where the deposited model and the ModelAngelo model agree or differ in amino acid identity. Residues with matching amino acid types between the two models generally exhibit higher pMDD values. In the enlarged segment shown in Figure 2e, the mismatch originates from a modeling error in the deposited PDB structure, where the residue register is shifted relative to the density. In this region, pMDD are reduced. Although pMDD is derived purely from the density map and is not intended as a dedicated map–model validation (Terashi *et al*., 2022), its ability to highlight locally unreliable map regions can still provide useful cues for identifying segments that warrant closer inspection during model building process. Importantly, because it does not require an input model, pMDD is available at an earlier stage of structure interpretation.

Building on the above case study, we further examine the relationship between pMDD and Q-score at both the per-map and global levels (see Supplementary Figure S6 for complementary analysis based on local resolution). As shown in Figure 3a, the distribution of per-map SCC between pMDD and Q-score shows a consistent trend across individual maps, with an average SCC of 0.611. We further compare the relationship between uncertainty and Q-score before and after calibration using 230,752 residues from 143 test maps (Figure 3b). While the uncalibrated MAE exhibits a weak association with Q-score, the calibrated pMDD shows a markedly clearer and more monotonic relationship. This result indicates that the proposed calibration effectively removes systematic biases and enables pMDD to serve as a consistent and interpretable indicator of local quality of enhanced-maps across maps.

**Figure 3:**
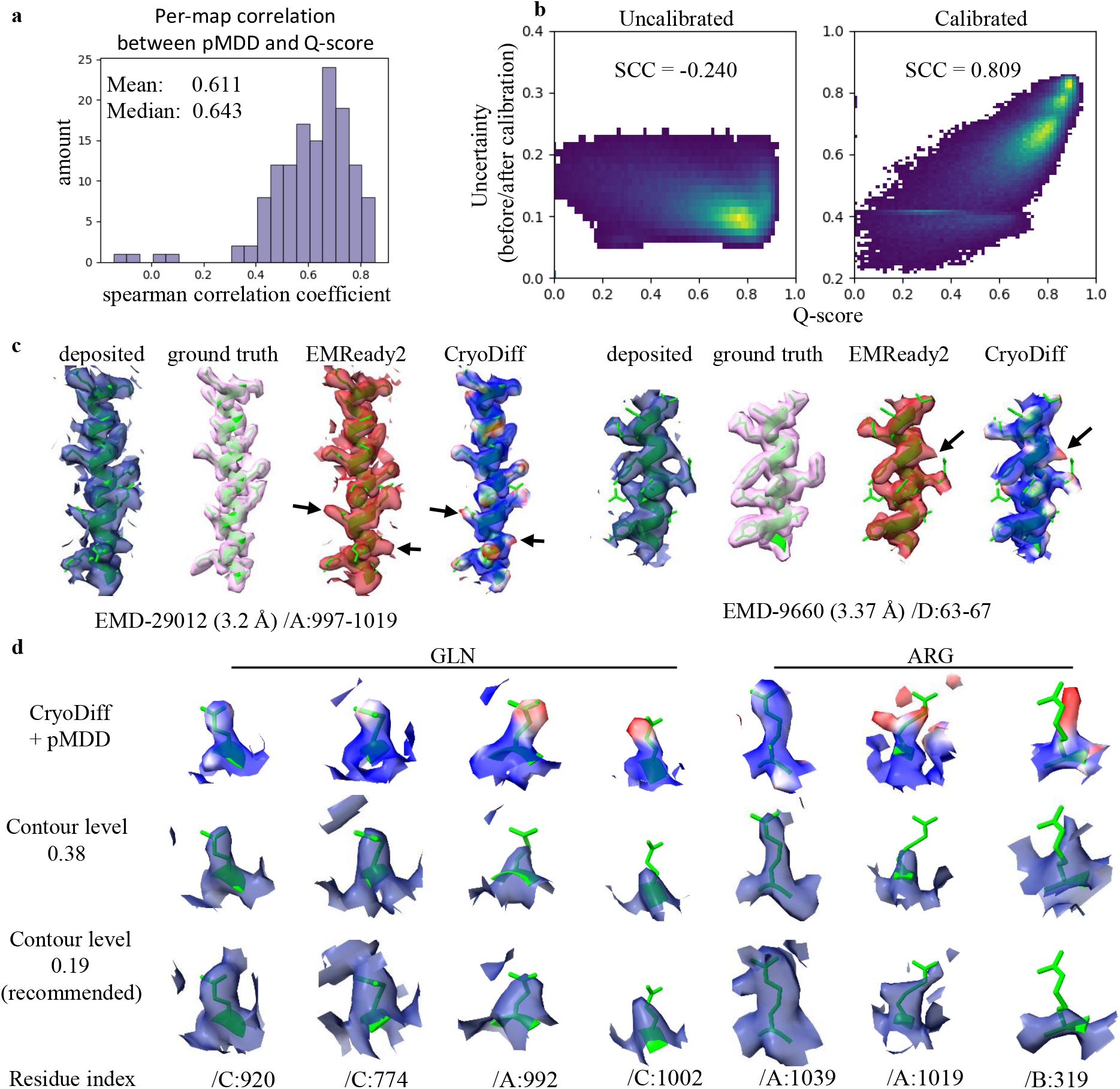
Extended evaluations on pMDD. **a** Distribution of per-map Spearman correlation coefficients (SCC) between Q-score and pMDD, showing the consistency of their association across individual maps. **b** Two-dimensional density plots comparing the relationship between uncertainty and Q-score before and after calibration across 230,752 residues. **c** Examples EMD-29012 (3.2 Å, PDB:8FDT) and EMD-9660 (3.37 Å, PDB:6IG0) that pMDD highlight artifact side chains generated by EMReady2 and CryoDiff. **d** Examples of side-chain density in EMD-21375 (3.46 Å, PDB:6VSB) with pMDD. The first row shows CryoDiff-processed maps colored by pMDD. The second and third rows display the deposited maps at different contour levels.

To further illustrate the practical relevance of pMDD, we present representative examples linking pMDD to local reconstruction artifacts and unreliable density features. Although the authors of EMReady have shown that their enhancement method is generally robust and does not introduce large-scale misleading structures, we observe that both EMReady2 and CryoDiff can occasionally introduce spurious side-chain densities at the local level, often appearing as side chains extending beyond the length supported by the deposited structure. These artifacts are likely associated with local over-sharpening of the input map or the misinterpretation of noise and reconstruction artifacts. In Figure 3c, cases from EMD-29012 (3.2 Å, PDB:8FDT) (Ton *et al*., 2023) and EMD-9660 (3.37 Å, PDB:6GI0) (You *et al*., 2019) are shown, where pMDD highlights side-chain features (black arrows) that likely correspond to such artifacts in the enhanced maps produced by EMReady2 and CryoDiff. In these regions, reduced pMDD values coincide with the anomalous density features, indicating that pMDD effectively identifies areas where side-chain interpretation may be unreliable. In Figure 3d, residues from EMD-21375 (3.46 Å, PDB:6VSB) (Wrapp *et al*., 2020) are arranged from left to right according to decreasing pMDD. The ordering is manually curated based on the map appearance at the chosen contour level for visualization purposes. The corresponding deposited maps exhibit a gradual decline in density clarity, transitioning from well-defined side-chain features to weaker or partially missing densities. Notably, an Arginine residue exhibiting density consistent with alternative conformations (rotameric variability) corresponds to a reduced pMDD value. These examples suggest that lower pMDD values are associated with reduced side-chain resolvability or increased conformational heterogeneity. During model building, atomic structures must balance chemical plausibility with agreement to the density map. In this context, pMDD provides a useful guide for this trade-off: regions with high pMDD typically correspond to well-resolved density and support confident modeling, whereas regions with low pMDD often reflect ambiguity in the density and may require further consideration.

### 2.3 Example results of the CryoDiff-processed maps

To quantify the performance of CryoDiff, we measured the map-model compatibility for five representative structures, with their corresponding PDB models as reference. As shown in Figure 4 and 5, we demonstrate that CryoDiff-processed maps consistently achieve superior compared to the deposited maps and EMReady2-processed maps.

**Figure 4:**
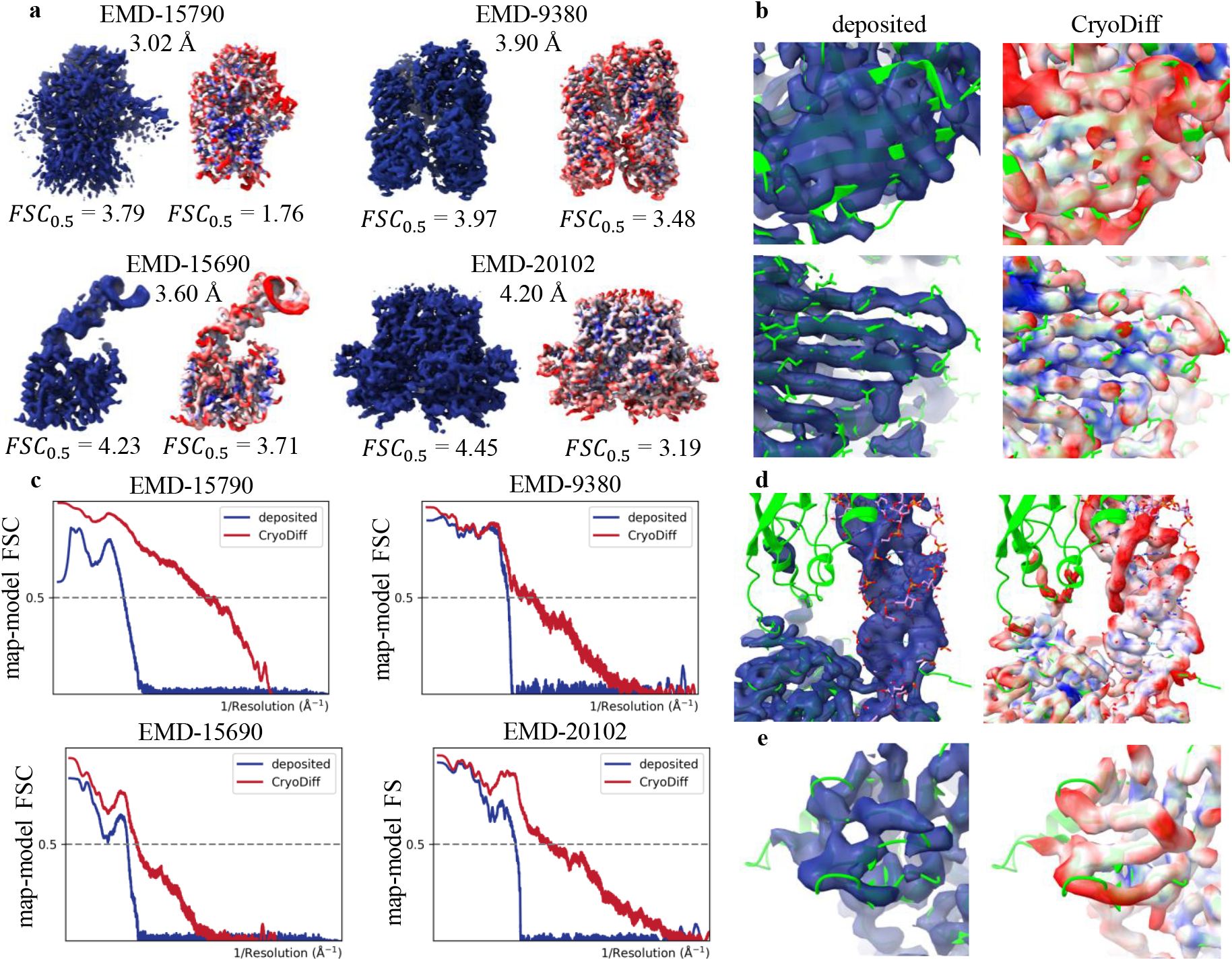
Four examples of CryoDiff-processed maps. **a** The deposited maps (left), and the CryoDiff-processed maps (right) which are colored by pMDD values. **b** Comparison between the deposited map and the CryoDiff-processed map over two *β*-sheet regions in truncated HIV-1 complex (EMD-9380, 3.90 Å), with the reference PDB model (PDB:6NIL) shown in green. **c** Map-model FSC curves for the deposited and CryoDiff-processed maps shown in **a. d** Comparison between the deposited map and the CryoDiff-processed map over a coil region in EMD-20102. **e** Comparison between the deposited map and the CryoDiff-processed map over a nucleic acids region in EMD-15690.

**Figure 5:**
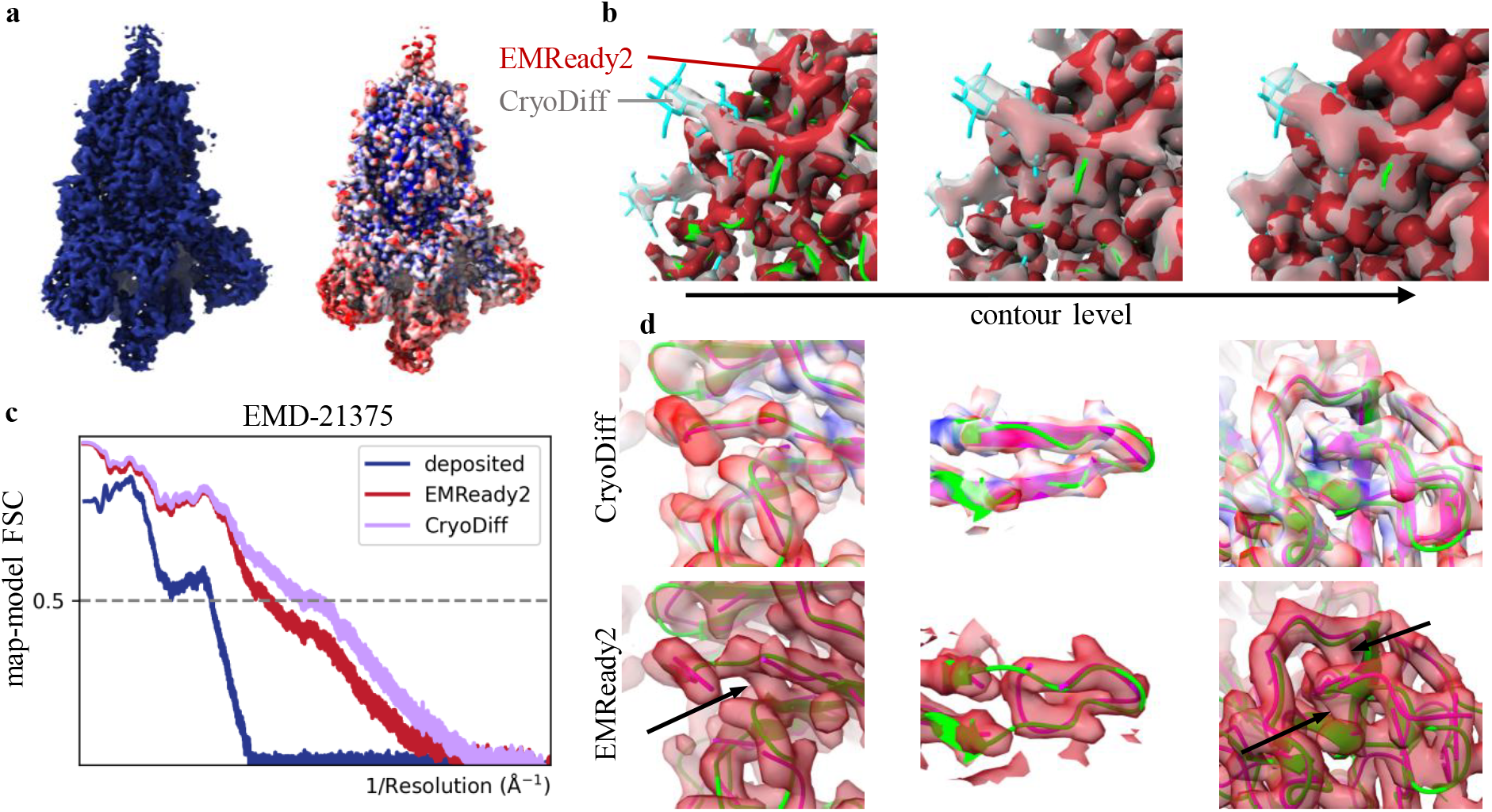
Improvement analysis for Prefusion 2019-nCoV spike glycoprotein (EMD-21375, 3.46 Å, PDB:6VSB). **a** The deposited map (left), and the CryoDiff-processed map (right) which is colored in pMDD values. **b** Comparison of local maps enhanced by EMReady2 (red) and CryoDiff (gray), displayed at various contour levels. **c** Map-model FSC curves for EMD-21375. **d** Comparison of the S1 subunit density from EMD-21375 enhanced by EMReady2 and CryoDiff. The reference PDB model is shown in green, and the *de novo* modeling result by ModelAngelo is shown in pink.

Figure 4a presents four CryoDiff-processed maps colored by pMDD valuse with the corresponding deposited maps on the left, including apolipoprotein N-acyltransferase Lnt from E.coli in complex with FP3 (Smithers *et al*., 2023) (EMD-15790, 3.02 Å, PDB:8B0O), truncated HIV-1 Vif/CBFbeta/A3F complex (Hu *et al*., 2019)(EMD-9380, 3.90 Å, 6NIL), Tb ADAT2/3 deaminase in complex with tRNA (Dolce *et al*., 2022)(EMD-15690, 3.60 Å, PDB:8AW3), and CDTb Pre-Insertion form Modeled (Anderson *et al*., 2020)(EMD-20102, 4.20 Å, PDB:6OKR). To quantify this comparison, the map-model FSC curves are presented in Figure 4c with their respective *FSC*_0.5_ values listed in Figure 4a.

Figure 4b demonstrates that pMDD values faithfully correlate with local map quality, using the deposited PDB model (PDB: 6NIL) as a reference. The upper panel highlights a *β*-sheet with poor initial resolution where the strands remain ambiguous even after CryoDiff processing. Our method correctly assigns low pMDD values to this area, signifying high uncertainty. In contrast, the lower panel showcases an adjacent, well-resolved *β*-sheet. Here, CryoDiff further enhances well-defined side-chain features, and the region is consistently assigned high pMDD values overall. Figure 4d and e show a coil region in EMD-20102 and a nucleic acids region in EMD-15690, respectively. These examples validate that CryoDiff significantly enhances the density within these regions, leading to better-connected main chains and more distinguishable nucleic acid backbones and bases.

We also demonstrate the superiority of CryoDiff over existing methods through a comprehensive case study of the prefusion 2019-nCoV spike glycoprotein (EMD-21375, 3.46 Å) (Wrapp *et al*., 2020) in Figure 5. Figure 5a presents the deposited map (left), and the CryoDiff-processed map (right) colored in pMDD value for EMD-21375. As presented in Figure 5c, the map-model FSC curves show that the CryoDiff-processed map achieves a resolution of 2.69 Å at *FSC*_0.5_, outperforming both the deposited map (4.47 Å) and EMReady2-processed map (3.18 Å). Figure 5b compares the density for N-acetyl-D-glucosamine (NAG) in maps processed by EMReady2 and CryoDiff at various contour levels. NAG is a critical component for stabilizing the spike protein and forming the glycan shield that modulates immune recognition (Huang *et al*., 2022). CryoDiff (gray) provides clearer density for the NAG residue than EMReady2 (red) across multiple contour levels, improving the visibility of glycan features that are essential for structural interpretation. Figure 5d demonstrates CryoDiff’s superior performance in improving backbone trace quality compared to EMReady2 within three regions of the S1 subunit domains. Specifically, in the EMReady2-processed map, the backbone density displays multiple incorrect connections and breaks (black arrows), leading to a fragmented *de novo* model (pink). In contrast, the CryoDiff-processed map yields a continuous and correctly connected backbone density that closely aligns with the reference PDB model (green), thereby enabling the generation of a more complete and accurate *de novo* model by ModelAngelo (build no seq).

### 2.4 Quantitative evaluation against other methods

To quantitatively validate the capability of CryoDiff in cryo-EM map enhancement, we calculated various model-map correlation metrics for both primary and half-map sets. Here, primary maps refer to the final post-processed maps deposited in the EMDB, while half-maps represent independent reconstructions from separate halves of the particle dataset and the average maps were used as input. We benchmarked CryoDiff against two state-of-the-art enhancement methods, CryoTEN and EMReady2, on two separate datasets comprising 143 primary maps and 80 half-map pairs. As presented in Table 1 and 2, CryoDiff consistently outperforms competing approaches across all evaluation metrics, including map-model *FSC*_0.5_, Q-score(P), Q-score(N), *CC*_*box*_, *CC*_*mask*_, and *CC*_*peaks*_.

**Table 1:** Map–model correlation of the deposited and processed maps on 143 primary maps.

| Method | $FS C_{0.5}$ | Q-score(P) | Q-score(N) | $CC_{box}$ | $CC_{mask}$ | $CC_{peaks}$ |
| --- | --- | --- | --- | --- | --- | --- |
| deposited | 4.981±3.14 | 0.531±0.15 | 0.466±0.15 | 0.719±0.10 | 0.774±0.09 | 0.626±0.11 |
| CryoTEN | 3.921±1.71 | 0.531±0.17 | 0.457±0.16 | 0.847±0.10 | 0.780±0.11 | 0.731±0.16 |
| EMReady2 | 3.401±1.83 | 0.589±0.17 | 0.507±0.17 | 0.854±0.07 | 0.791±0.10 | 0.745±0.14 |
| **CryoDiff** | **3.045±1.75** | **0.608±0.17** | **0.523±0.17** | **0.868±0.07** | **0.816±0.07** | **0.778±0.11** |

All values are reported as mean ± standard deviation across the test dataset.

**Table 2:**
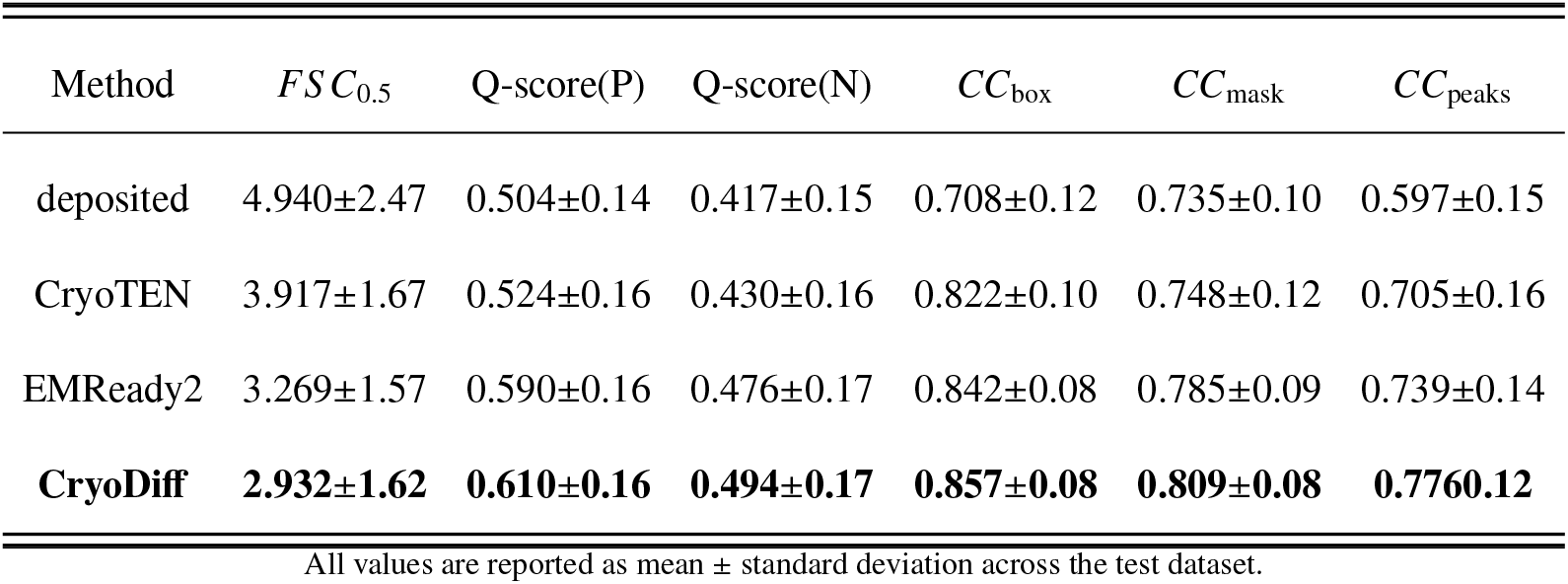
Map-model correlation of the deposited and processed maps on 80 pairs of half maps.

| Method | $FS C_{0.5}$ | Q-score(P) | Q-score(N) | $CC_{\text{box}}$ | $CC_{\text{mask}}$ | $CC_{\text{peaks}}$ |
| --- | --- | --- | --- | --- | --- | --- |
| deposited | 4.940±2.47 | 0.504±0.14 | 0.417±0.15 | 0.708±0.12 | 0.735±0.10 | 0.597±0.15 |
| CryoTEN | 3.917±1.67 | 0.524±0.16 | 0.430±0.16 | 0.822±0.10 | 0.748±0.12 | 0.705±0.16 |
| EMReady2 | 3.269±1.57 | 0.590±0.16 | 0.476±0.17 | 0.842±0.08 | 0.785±0.09 | 0.739±0.14 |
| **CryoDiff** | **2.932±1.62** | **0.610±0.16** | **0.494±0.17** | **0.857±0.08** | **0.809±0.08** | **0.7760.12** |

All values are reported as mean ± standard deviation across the test dataset.

In terms of map-model *FSC*_0.5_ shown in Table 1 and 2, CryoDiff achieved an average value of 3.044 Å across the primary map set, representing a significant improvement over the deposited maps (by 1.937 Å), CryoTEN (by 0.877 Å), and EMReady2 (by 0.356 Å). Furthermore, CryoDiff attained a median *FSC*_0.5_ of 2.42 Å, surpassing CryoTEN and EMReady2 by 1.07 Å and 0.63 Å, respectively. For the half-map set, CryoDiff yielded an average *FSC*_0.5_ value of 2.932 Å, showing consistent improvement over all baselines. Specifically, CryoDiff surpassed the deposited maps by 2.931 Å, CryoTEN by 0.985 Å, and EMReady2 by 0.337 Å. Additionally, CryoDiff achieved a median map-model *FSC*_0.5_ of 2.26 Å, surpassing CryoTEN and EMReady2 by 1.28 Å and 0.70 Å, respectively. These improvements in map–model *FSC*_0.5_ values indicate that CryoDiff recovers richer high-frequency details and yields better-resolved structural features, demonstrating its robustness across different datasets.

We then employed the Q-score metric to measure atom resolvability in Cryo-EM maps (Pintilie *et al*., 2020). As shown in Table 1 and 2, CryoDiff successfully improves the Q-score for both proteins and nucleic acids. Specifically, on the primary map set, the average Q-score improved significantly from 0.531 to 0.608 for proteins (Q-score(P)) and from 0.466 to 0.523 for nucleic acids (Q-score(N)). Similarly, for the half-map set, the average Q-score(P) increased from 0.504 to 0.610, while the average Q-score(N) rose from 0.417 to 0.494. These enhancements suggest that CryoDiff effectively recovers density features, leading to more interpretable maps for both proteins and nucleic acids.

We also report the real-space correlation values between model-derived maps and enhanced maps (Afonine *et al*., 2018) in Table 1 and 2. Three distinct correlation coefficient metrics were used to evaluate CryoDiff’s performance, including *CC*_box_, *CC*_mask_, and *CC*_peaks_. CryoDiff demonstrated substantial improvements across all these metrics. On the primary map set, the mean *CC*_box_, *CC*_mask_ and *CC*_peaks_ of the CryoDiff processed maps reached 0.868, 0.816 and 0.778 respectively, representing significant improvements over the deposited maps of 0.719, 0.774, and 0.626. Similarly, on the half-map set, CryoDiff achieves 0.857, 0.809 and 0.776 for *CC*_box_, *CC*_mask_ and *CC*_peaks_, respectively. These results also reflect a substantial improvement over the deposited maps of 0.708, 0.735, and 0.597, respectively.

These results demonstrate that CryoDiff improves the quality of both primary and half maps across all metrics, with slightly greater gains observed for primary maps (see Supplementary Figure S1).

### 2.5 Improvement in map interpretability

The ultimate goal of density modification is to facilitate the construction of atomic models. To quantify how map enhancement improves the accuracy of downstream *de novo* atomic modeling, we evaluated models built from maps processed by CryoDiff and other methods based on residue coverage and sequence matching results.

Specifically, we performed chain-wise model building using the traditional automated tool phenix.map to model. And there are 651 protein chains that were first extracted from 143 primary density maps using phenix.map box. phenix.map to model was then employed to automatically build atomic structures for these extracted chains. Finally, the built chains were compared with their corresponding deposited structures from the PDB using phenix.chain comparison to obtain the evaluation metrics.

As illustrated in Table 3, CryoDiff-processed maps consistently yielded superior atomic models. CryoDiff achieved an average residue coverage of 77.0% and sequence matching of 48.6%. This is significantly higher than the 59.9% residue coverage and 31.7% sequence matching obtained from the deposited maps. Furthermore, CryoDiff consistently outperformed other benchmark methods, including CryoTEN (64.3% residue coverage, 35.1% sequence matching) and EMReady2 (74.7% residue coverage, 45.2% sequence matching). This superior performance demonstrates CryoDiff’s effectiveness in improving the interpretability of cryo-EM density maps for better automated model building.

**Table 3:** Comparison of map interpretability by phenix.map to model.

|  | Residue coverage(%) | Sequence matching(%) |
| --- | --- | --- |
| deposited | 59.943±24.77 | 31.705±26.92 |
| CryoTEN | 64.311±24.3 | 35.149±29.15 |
| EMReady2 | 74.712±20.6 | 45.203±30.7 |
| **CryoDiff** | **77.017±18.99** | **48.562±31.59** |

All values are reported as mean ± standard deviation across the test dataset.

We further demonstrate that CryoDiff-processed maps substantially improve the completeness and accuracy of atomic models generated by deep-learning-based methods (Figure 6). Specifically, we employed ModelAngelo, a state-of-the-art *de novo* modeling method. To adapt ModelAngelo to the density characteristics of CryoDiff-processed maps, we first fine-tuned its amino-acid type prediction module. We then applied this fine-tuned model to a test set of 127 cryo-EM maps. As shown in Figure 6a, models built from CryoDiff-processed maps exhibit higher residue coverage while maintaining comparable precision relative to those built from deposited maps and EMReady2-processed maps. On average, residue coverage increased from 76.0% for deposited maps to 77.4% for EMReady2-processed maps and further to 81.6% for CryoDiff-processed maps, while backbone precision remained largely unchanged (98.4%, 98.4%, and 98.2%, respectively). We also assessed model completeness, a metric integrating both backbone recall and amino acid accuracy. As shown in Figure 6b,CryoDiff consistently improved performance across most test cases, increasing the mean completeness from 73.8% for deposited maps to 75.5% for EMReady2-processed maps and further to 79.3% for CryoDiff-processed maps. In terms of sequence identification presented in Figure 6c, models derived from CryoDiff-processed maps achieved a mean amino acid accuracy of 96.7%, comparable to the 96.8% obtained with EMReady2-processed maps and slightly higher than the 95.9% obtained with deposited maps. Furthermore, models built from processed maps also exhibited improved geometric accuracy. The C*α* RMSD decreased from 0.632 Å for deposited maps to 0.612 Å for EMReady2-processed maps and remained comparable at 0.616 Å for CryoDiff-processed maps, while backbone RMSD was reduced from 0.710 Å to 0.695 Å and 0.705 Å, respectively. Beyond the high-confidence pruned structures, we also analyzed the raw output structures, which are critical for identifying ambiguous regions and guiding manual refinement. Full evaluations for both structure types are detailed in Supplementary Table 3. Representative examples are shown in Figure 6d. For both the glucagon receptor bound to glucagon and *β*-arrestin 1 (Chen *et al*., 2023) (EMD-36607, 3.3 Å, PDB: 8JRV) and the SERCA2b (Zhang *et al*., 2022) (EMD-32347, 3.4 Å, PDB: 7W7T), models built from CryoDiff-processed maps exhibit improved completeness relative to those derived from deposited and EMReady2-processed maps.

**Figure 6:**
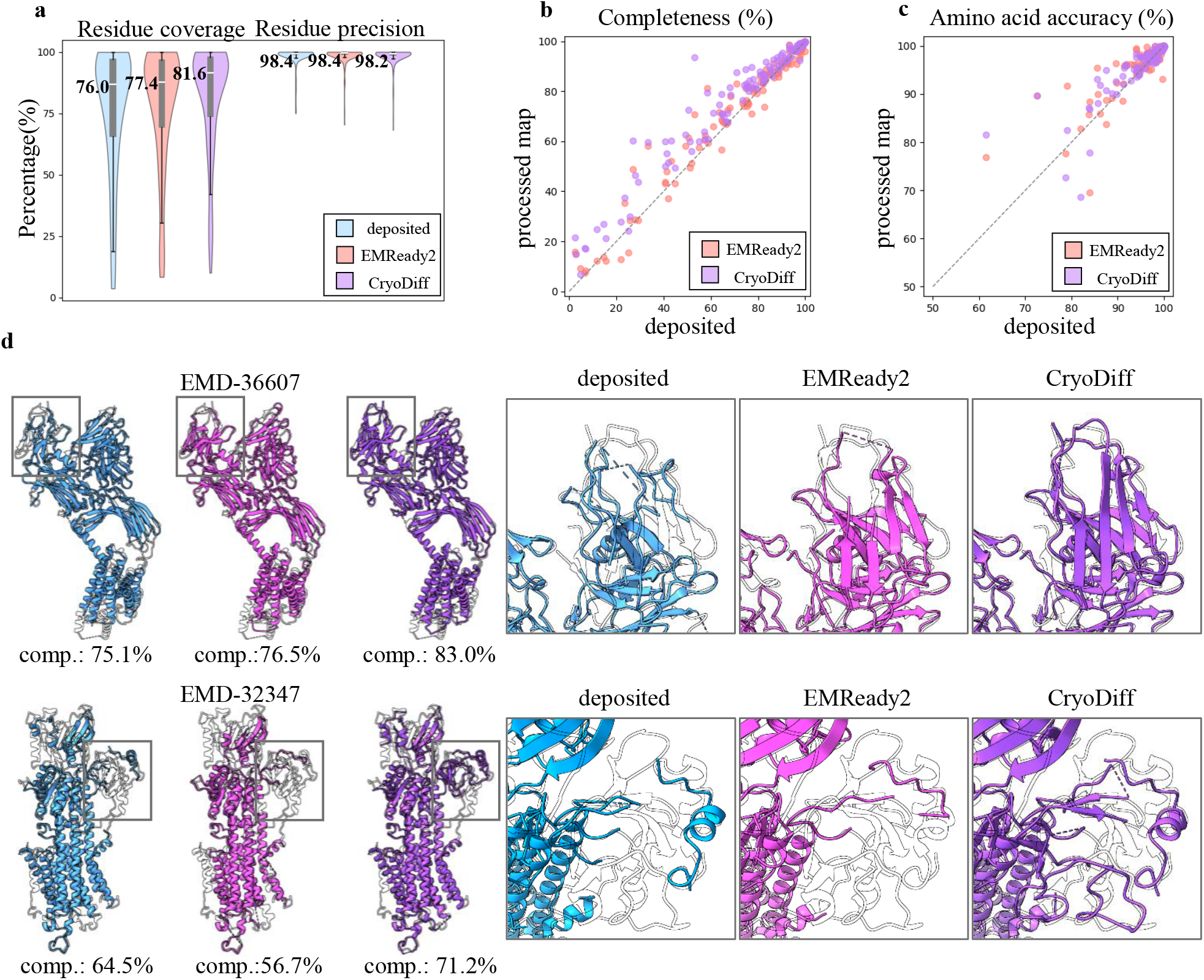
Improved ModelAngelo-built models via CryoDiff. **a** Violin plot comparing the backbone recall and precision ModelAngelo-built models using deposited, EMReady2-processed and CryoDiff-processed maps. **b** Scatter plots comparing the completeness of ModelAngelo-built models using deposited EMReady2-processed and CryoDiff-processed maps. **c** Scatter plots comparing the amino acid accuracy of ModelAngelo-built models using deposited EMReady2-proceesed and CryoDiff-processed maps. **d** Examples of *de novo* models built from the deposited (blue, left), EMReady2-processed (pink, middle), and CryoDiff-processed maps (purple, right). PDB models are shown as silhouettes. Enlarged regions for both examples are shown on the far right.

## 3 Method

### 3.1 Data collection

To establish a benchmark dataset for model training and evaluation, we curated a collection of cryo-EM map-model pairs from the EMDB and PDB databases, restricting to entries with resolution better than 7 Å. To ensure a high standard of data quality and relevance, we implemented two critical data filtering criteria. First, maps exhibiting a model–map correlation coefficient *CC*_box_ below 0.5 were excluded. Second, to further reduce redundancy, we utilized mmseqs.easy-linclust (Hauser *et al*., 2016) to cluster the remaining entries with a chain-level sequence identity threshold of 30%. This process finally yielded 980 non-redundant entries, which were randomly partitioned into training (737), validation (100), and testing (143) sets. For the test data, all 143 entries constitute the primary test set, while a subset of 80 entries with publicly available half-maps forms the half-map test set. Notably, complexes containing RNA components were explicitly retained. The final dataset encompasses a resolution range of 1.8 Å to 6.8 Å, covering both high- and medium-resolution regimes (see Supplementary Figure S5). All entries included in this study were collected up to May 2025.

### 3.2 Data preprocessing

To ensure data consistency, the experimental maps are first resampled to a uniform voxel size of 1 Å through cubic interpolation. Subsequently, negative densities are truncated to zero, and the volumes are normalized to the [0,1] scaled against the 99.999th percentile. Ground-truth simulated maps *ρ*(**x**) are then computed from the corresponding atomic models on the aligned experimental grids **x** using the molmap command in UCSF ChimeraX (Pettersen *et al*., 2004), which can be formulated as:

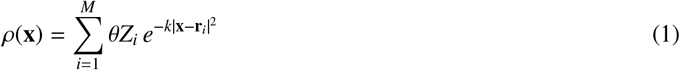

where *Z*_*i*_ and *r*_*i*_ is the atomic number and the position vector of the i-th heavy atom among the total of *M* heavy atoms, *k* = (*π/*1.2 + 0.6*R*)^2^ with *R* is the reported resolution of the experimental maps and the scaling factor *θ* is defined as *θ* = (*k/π*)^1.5^. Finally, these aligned map pairs were first cropped into boxes of the size 60^3^ with a stride of 30, followed by a random crop to a final size of 48^3^ for model training. Boxes containing no positive values were excluded.

### 3.3 Network architecture

We employed a conditional diffusion model (Rombach *et al*., 2022b) to enhance cryo-EM density maps, leveraging its ability to overcome the blurring effects of conventional denoising and recover fine structural details. Inspired by LDM (Rombach *et al*., 2022a), we adopted its time-conditional 2D U-Net backbone and extended it into time- and map-conditional 3D U-Net architecture to serve as our reverse denoising network. The overview of CryoDiff is presented in Figure 1. This 3D U-Net network consists of a hierarchical architecture comprising four encoder stages, a central bottleneck, and four decoder stages, bridged by skip connections. To balance feature extraction capabilities, residual convolutional blocks are applied across all scales for local modeling, whereas global attention blocks are selectively integrated at deeper stages to capture long-range dependencies between the noisy input and the conditioning cryo-EM densities. The detail of the architecture is shown in Supplementary Figure S4.

### 3.4 Network training

CryoDiff is built upon a denoising diffusion framework that involves a fixed forward process that progressively adds noise to the original signal, and a learned reverse process. This reverse process is trained to iteratively recover the target signal, guided by the diffusion time step *t*, and other conditional information. For the map enhancement task, CryoDiff conditions on the experimental box *x*_*c*_ to recover the high-resolution signal, represented by the simulated box *x*_0_.

The forward process generates a noisy sample *x*_*t*_ from the clean data *x*_0_ at the given timestep *t*. This is achieved by sampling a noise variable *ϵ* from a standard Gaussian distribution, *ϵ* ∼ 𝒩(0, *I*), with the same dimensionality as *x*_0_. Thus, the *x*_*t*_ can be linearly expressed as:

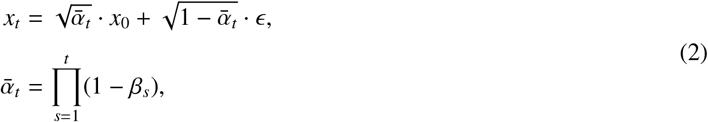

where *β*_*t*_ is the noise schedule increasing linearly from *β*_*start*_ = 0.00085 to *β*_*end*_ = 0.025 over *T* diffusion timesteps (*T* = 1000).

The reverse process aims to predict the noise component 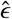 embedded within the noisy box *x*_*t*_ at an arbitrary timestep *t*. It is performed by a 3D U-Net model, denoted as **G**(·), which is conditioned on the noisy simulated box *x*_*t*_, experimental box *x*_*c*_ and random timestep *t*, which can be formulated as:

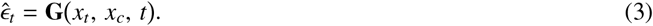

Inspired by EMReady, we employ a hybrid loss function ℒ_**G**_, which is the sum of Smooth L1 loss ℒ_*smooth*_ and the SSIM loss ℒ_*ssim*_, to calculate the difference between ground-truth noise *ϵ* and the predicted noise 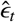 at an arbitrary timestep *t*.

The Smooth L1 loss ℒ_*smooth*_ focuses on local difference of *ϵ* and 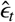, and is formulated as:

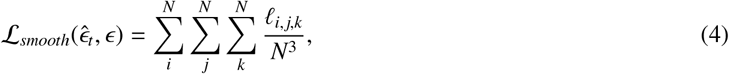

where *N* is the side length of the output box (*N* = 48) and *ℓ*_*i, j*,*k*_ is the Smooth L1 distance between ground-truth *ϵ* and predicted 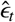 at position (*i, j, k*):

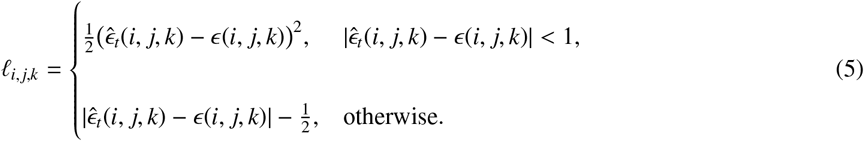

The SSIM loss ℒ_*ssim*_ calculates the global difference of *ϵ* and 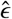, and is formulated as:

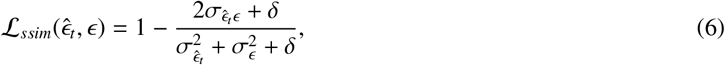

where *δ* = 10^−6^, 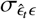 denotes the covariance between 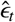 and 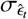, and *σ*_*ϵ*_ represent the standard deviations of 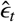 and *ϵ*, respectively.

### 3.5 Conditional enhanced map inference

The conditional generation of the enhanced map is guided by the input experimental map and performed via a multi-stage inference pipeline. First, the experimental map is decomposed into overlapping boxes of size 48^3^ with a stride of 16. To improve computational efficiency, boxes are subsequently excluded from processing if their maximum voxel value does not exceed 3*σ*, where *σ* is the root-mean-square value of the experimental map. For each remaining box, the enhanced result is then generated using a denoising diffusion implicit model (DDIM) sampler (Song *et al*., 2020). To accelerate inference, the sampler operates on a sparse subsequence *τ* = {*τ*_1_, …, *τ*_*S*_} of *S* timesteps (*S* = 20). This subsequence *τ* is constructed by uniformly sampling from the original *T* timesteps of the forward process. The reverse diffusion process **R**(·) begins with the random noise 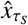 sampled from a Gaussian distribution and is conditioned on the experimental box *x*_*c*_, which can be formulated as:

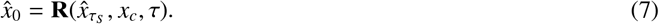

This process is executed iteratively over the subsequence *τ*, and can be formulated as:

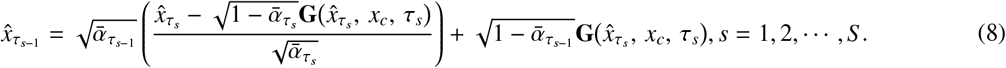

Finally, all resulting enhanced boxes 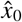 are assembled into the full-sized enhanced map by averaging the overlapping region.

### 3.6 Local confidence assessment with pMDD

A comprehensive post-processing workflow should not only produce high-quality enhanced density maps but also provide quantitative assessments of their reliability, which are critical for guiding subsequent model building and validating structural interpretations. To address this need, we introduce a novel voxel-wise metric, the predicted Multi-Diffusion samples Difference (pMDD). This metric is specifically designed to leverage the inherent uncertainty of the reverse diffusion process as an indicator for model confidence, thereby offering a direct measure of local map quality, which is further refined through an uncertainty calibration procedure to improve its interpretability.

This metric measures the local consistency and stability of the inference results by calculating the mean absolute error (MAE) between the multiple uncertainty boxes and the enhanced box 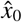. Specifically, CryoDiff first utilizes its reverse process to generate a high-quality assembly result, which serves as both the reference map for comparison and the final enhanced map. Next, the original experimental map and the final enhanced map are partitioned into overlapping boxes *x*_*c*_ and 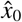 of size 48^3^ using a stride of 24. For each experimental box *x*_*c*_, it then generates another *M* (*M* = 5, see Supplementary Figure S9) inferred boxes 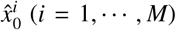 based on *M* different random noise 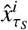 sampled from a Gaussian distribution. The initial (uncalibrated) pMDD is defined as:

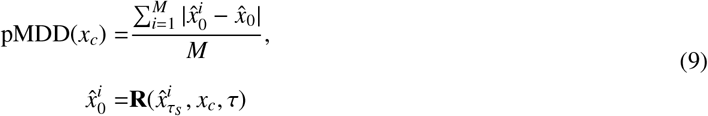

To improve the comparability and interpretability of pMDD across maps with different global resolutions, we further perform an uncertainty calibration procedure. Specifically, residue-wise (uncalibrated) pMDD are first computed and paired with local Q-score, which serve as a reference for structural resolvability. To account for the influence of global resolution, the dataset is partitioned into multiple resolution bins. Within each bin, isotonic regression is applied to learn a monotonic mapping from raw pMDD to the corresponding Q-score. The learned mappings (see Supplementary Figure S8) are then used to transform raw pMDD values into calibrated score. This calibration yields the final normalized and resolution-consistent pMDD to be comparable across datasets while preserving the relative ordering of voxel-wise MAE.

### 3.7 Evaluation metrics

We utilize various model-map corelation metrics including map-model *FSC*_0.5_, Q-score, *CC*_box_, *CC*_mask_ and *CC*_peaks_ to verify the superiority of CryoDiff.

#### Map-model *FSC*_0.5_

The map-model Fourier Shell Correlation (*FSC*) provides a quantitative measure of similarity between the model-derived map and the enhanced map as a function of spatial frequency, with the resolution of this match conventionally defined by the point where the FSC curve intersects the 0.5 threshold (Böttcher *et al*., 1997).

#### Q-score

Q-score (Pintilie *et al*., 2020) is a quantitative metric that characterizes the resolvability of individual atoms in cryo-EM maps. It is defined as the correlation coefficient between two vectors, denoted as **u** and **v**, which represent the density values sampled from the input map and the corresponding simulated map in the local neighborhood of a given atom, respectively:

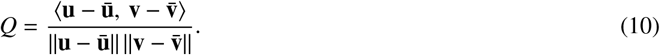

Once per-atom Q-score were computed, we determined the average Q-score(P) and Q-score(N) values separately for all protein atoms and for all nucleic acid atoms within the entry. And the Q-score is calculated by the MapQ plugin in UCSF Chimera (Pettersen *et al*., 2004).

#### CC calculation

The map-model real-space cross-correlation (CC) (Afonine *et al*., 2018) aims to quantify how well the model fits the experimental map. It is calculated between the experimental (enhanced) map *ρ*_1_(**i**) and a model-derived map *ρ*_2_(**i**), which is first generated from the atomic coordinates and then sampled onto the identical grid **i** as the experimental map. The process of CC calculation can be formulated as:

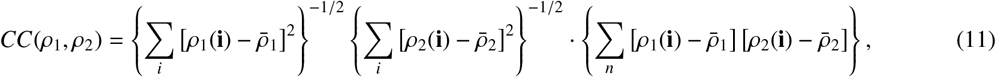

where 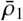 and 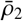 is the mean value of *ρ*_1_, *ρ*_2_. In this work, we utilize *CC*_box_, *CC*_mask_, and *CC*_peaks_ to quantify the quality of the enhanced map with phenix.map model cc tool. Specifically, *CC*_box_ assesses global map completeness, *CC*_mask_ focuses on local model accuracy, and *CC*_peaks_ targets the robust consistency of the strongest signal peaks. Each metric is calculated separately from the filtered grid **n** based on its specific target.

#### Residue coverage

Residue coverage (Terashi and Kihara, 2018) is used to evaluate the improvement in modeling accuracy provided by an enhanced map. Specifically, it quantifies the percentage of residues in the reference model whose C*α* atoms are located within 3 Å of their corresponding C*α* atoms in the built model.

#### Sequence matching

Sequence matching is used to evaluate the accuracy of amino acid type assignment in the built model. Specifically, it quantifies the percentage of matched residues whose amino acid types in the built model are identical to those in the reference model.

#### Residue precision

In contrast to residue coverage, residue precision is used to assess whether post-processing introduces structural artifacts during the model-building process. It quantifies the percentage of residues in the built model that have a matching C*α* atom within 3 Å of the reference model.

#### Completeness

Completeness integrates backbone recall with amino acid assignment accuracy. First, the Hungarian algorithm is employed to establish optimal one-to-one correspondences between the built and reference models. Let *n* denote the total number of residues in the reference model, and *p* denote the number of matched residue pairs with identical amino acid types. Completeness is thus defined as the fraction of reference residues that are correctly recovered in both position and type (i.e.,*p/n*).

#### Amino acid accuracy

Amino acid accuracy is used to quantify the correctness of amino acid type assignment in the built model relative to the reference model. It is computed over all matched residues (i.e. *p/m*). This metric measures the consistency of residue type prediction between the reconstructed and reference structures.

#### C*α* RMSD

The C*α* root-mean-square deviation (RMSD) measures the geometric accuracy of the built model relative to the reference model, and can be calculated as:

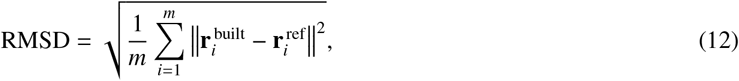

where 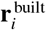 and 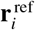 denote the coordinates of the C*α* atoms of the *i*-th residue in the built model and the reference model, respectively.

### 3.8 Network implementation

The proposed network was implemented in PyTorch 1.13.1 (CUDA 12.2) and trained on two NVIDIA A100 GPUs. To mitigate overfitting and enhance model generalization, data augmentation techniques, including random cropping and rotation, were applied during training. The network was optimized using the AdamW optimizer with a batch size of 32 and an initial learning rate of 10^−4^. The learning rate was dynamically reduced by 10% upon a validation loss plateau of six epochs to ensure stable convergence. The model was totally trained for 300 epochs.

### 3.9 The calculation of per-residue pMDD

To investigate the relationship between pMDD and Q-score and modeling error, we derived the residue-wise pMDD for a given structure. For Figure 2c and 3a, the pMDD for a residue is defined as the mean value within a mask obtained by intersecting a 5 Å sphere centered at the C*α* atom with a molecular mask from the enhanced map (contour level 0.1). We further evaluated the robustness of this definition using different sphere radii (see Supplementary Figure S7). For Figure 2d, the pMDD for a residue is defined as the mean value within a mask obtained as the union of 2 Å spheres centered at all backbone atoms.

## 4 Discussion

In this work, we proposed a diffusion-based model named CryoDiff for Cryo-EM map post-processing. The evaluation results demonstrate that CryoDiff consistently outperforms existing methods across multiple metrics. Specifically, maps processed by CryoDiff exhibit superior backbone continuity and better-resolved side-chain density features, thereby directly facilitating more accurate downstream structural modeling. The observed improvements underscore the intrinsic advantages of diffusion-based modeling. While conventional feed-forward methods often suffer from over-smoothing and the generation of artifactual features, CryoDiff overcomes these limitations by leveraging the iterative and probabilistic sampling inherent in diffusion processes. This allows the model to progressively reconstruct plausible density distributions conditioned on the experimental data. Consequently, this generative mechanism effectively mitigates over-smoothing and reduces the risk of artifacts, providing a more faithful representation of the underlying biological structures.

Furthermore, CryoDiff provides a voxel-wise reliability metric, termed pMDD, which is derived from the mean absolute error (MAE) across multiple diffusion samples and subsequently calibrated to correct for resolution-dependent biases, enabling a more interpretable measure of local resolvability. While we have demonstrated its efficacy in highlighting regions susceptible to misinterpretation and its correlation with local quality metrics such as Q-score, certain limitation should be considered. Namely, pMDD is influenced by multiple factors—including intrinsic structural flexibility, local resolution, and training data representation—rather than a single source of uncertainty. Consequently, a high pMDD value indicates general uncertainty but does not determine its specific cause.

Despite its superior performance, CryoDiff presents certain limitations. First, the model may exhibit limited efficacy in resolving features underrepresented in the current training data, such as small ligands, glycosylation sites, and metal-coordination spheres (see Supplementary Figure S2a). Additionally, CryoDiff may occasionally misidentify extremely weak density signals as noise, potentially leading to the loss of real structural information (see Supplementary Figure S2b). To mitigate this risk, manual inspection and cross-validation against the raw experimental map remain essential. Finally, the current implementation is computationally intensive (see Supplementary Figure S3). Thus, our future work will focus on optimizing inference strategies and exploring diffusion distillation techniques to enhance computational efficiency for large-scale applications.

## 5 Availability of data and materials

The data that support this study are available from the corresponding authors upon request. The data used as examples are available from the EMDB and PDB. The source data underlying Table 1, 2, 3 and Supplementary Tables S1 are provided as Source Data files.

## 6 Funding

This research was supported by the National Key Research and Development Program of China [2021YFF0704300], the Fundamental Research Funds for the Central Universities, the Dubai Future Foundation [Award No. 2024CANAD-MES-061], and the Natural Science Foundation of Ningxia Province [2026AAC040016], the National Natural Science Foundation of China [W2511070, 624B2089, 32241027, 62472034], and the Natural Science Foundation of Shandong Province [ZR2023YQ057].

## Notes

### Competing Interest Statement

The authors have declared no competing interest.

