## Supplementary material for "CryoDiff: An uncertainty-aware diffusion model for Cryo-EM map enhancement": Supplymentary Figure S1-S9, Table S1

### 1 Evaluation results

In this study, we use primary maps as training data, which refer to the final post-processed Cryo-EM reconstructions. In addition to this setting, evaluations of CryoDiff on raw half-maps also demonstrate notable improvements compared with the deposited raw half-maps, further indicating the generalization capability of CryoDiff. We therefore conduct a comparison using primary maps and half-maps as inputs to systematically assess their impact on CryoDiff performance. The results reveal that using a processed map can yield better results compared to using raw map. The results indicate that processed maps generally yield improved performance compared with raw maps. Therefore, applying B-factor-based sharpening to raw maps prior to CryoDiff inference can be beneficial.

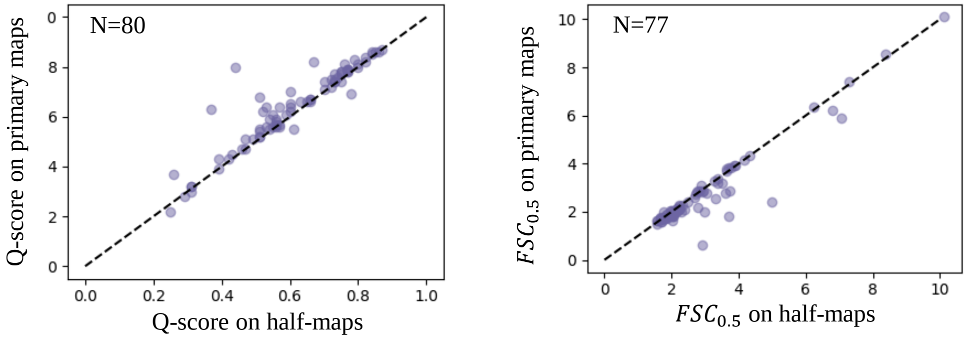


Figure S1: Comparison of Q-score and map-model $FSC_{0.5}$ between using primary maps and half-maps as input for CryoDiff.

Supplementary Table S1 summarizes ModelAngelo model-building performance across different map types. Here, coverage denotes residue coverage, precision denotes residue precision, compl. denotes completeness, and acc. denotes amino acid accuracy. C$\alpha$ RMSD and BB RMSD represent the root-mean-square deviations of C$\alpha$ atoms and backbone atoms, respectively. CryoDiff-processed maps consistently improve residue coverage and completeness for both pruned and raw structures compared with deposited and EMReady2-processed maps. Notably, these gains are achieved without sacrificing residue precision or geometric accuracy, as reflected by comparable C$\alpha$ and backbone RMSD values. In addition, the improvements are more pronounced in raw structures, indicating that CryoDiff provides more informative initial models for identifying uncertain regions and guiding subsequent refinement.

Table S1: Comparison of ModelAngelo model-building performance for both pruned and raw structure using deposited maps, EMReady2-processed maps, and CryoDiff-processed maps.

| Structure | Map type | Coverage | Precision | Compl. | Acc. | C$\alpha$ RMSD | BB RMSD |
|---|---|---|---|---|---|---|---|
| Pruned | Deposited map | 76.0 | 98.4 | 73.8 | 95.9 | 0.632 | 0.710 |
| | EMReady2-porcessed map | 77.4 | **98.4** | 75.5 | **96.8** | **0.612** | **0.695** |
| | CryoDiff-processed map | **81.6** | 98.2 | **79.3** | 96.7 | 0.616 | 0.705 |
| Raw | Deposited map | 95.0 | **91.4** | 76.8 | 80.3 | 0.776 | 0.809 |
| | EMReady2-processed map | 95.4 | 90.0 | 78.1 | 81.6 | 0.763 | 0.987 |
| | CryoDiff-processed map | **96.3** | 90.6 | **81.3** | **84.4** | **0.749** | **0.966** |

Taken together, these results indicate that CryoDiff-processed maps consistently improve map interpretability and downstream model-building performance across different tools and evaluation metrics. The observed gains in residue coverage and completeness suggest that CryoDiff enhances structurally informative features in the density maps. One plausible explanation is that CryoDiff produces more distinguishable side-chain densities, which can serve as reliable anchors for model-building algorithms. These anchors facilitate more accurate sequence registration and enable continuous backbone tracing, ultimately leading to more complete and accurate atomic models.

Figure S2 presents representative cases illustrating the potential limitations of CryoDiff enhancement results. In Figure S2a, several ligand-containing regions are shown where CryoDiff does not improve ligand density interpretability, including Chlorophyll *a* in EMD-6930 (PDB: 5ZGH) (Pi *et al.*, 2018) and MC3 in EMD-38322 (PDB: 8XGG) (Cho *et al.*, 2024). In these cases, ligand-associated densities remain weak or indistinct in the CryoDiff-processed maps, suggesting that ligand placement should be verified against the original deposited maps.

Figure S2b shows a linker region of a membrane protein in EMD-31487 (3.81Å, PDB:7F7F) (Xu *et al.*, 2022), where the original map contains extremely weak signal. Consequently, the CryoDiff-processed map exhibits no detectable density in the corresponding region.

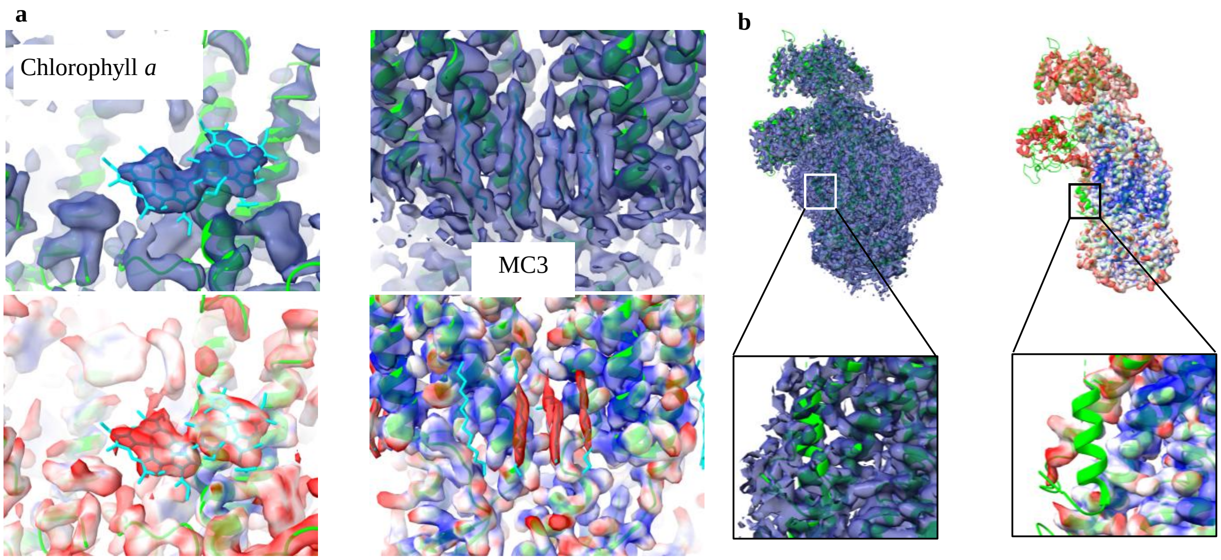


Figure S2: **a** Performance of CryoDiff on ligands, including Chlorophyll *a* (CLA) in EMD-6930 (PDB:5ZGH) and 1,2-dimyristoyl-rac-glycero-3-phosphocholine (MC3) in EMD-38322 (PDB:8XGG). **b** An example in EMD-31487 where the extremely weak signal in the original map leads to an absence of density in the corresponding region of the CryoDiff-processed map.

The left panel of Figure S3 shows the distribution of residue counts in the 143 test maps while the right panel shows the distribution of CryoDiff inference time, including both enhanced map generation and uncertainty estimation, Notably, approximately 80% of the test maps can be processed within 1000 seconds.

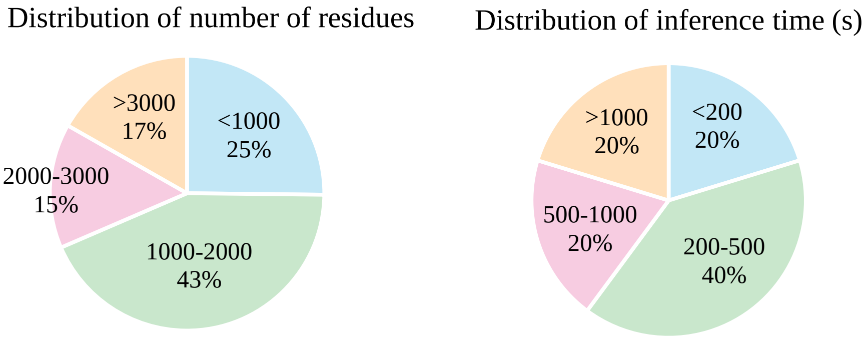


Figure S3: Left panel shows the distribution of numbers of residues in the 143 test maps and the right panel shows the inference time (including the generation of the enhanced map and pMDD map) distribution of CryoDiff.

#### 2 Details of model architecture

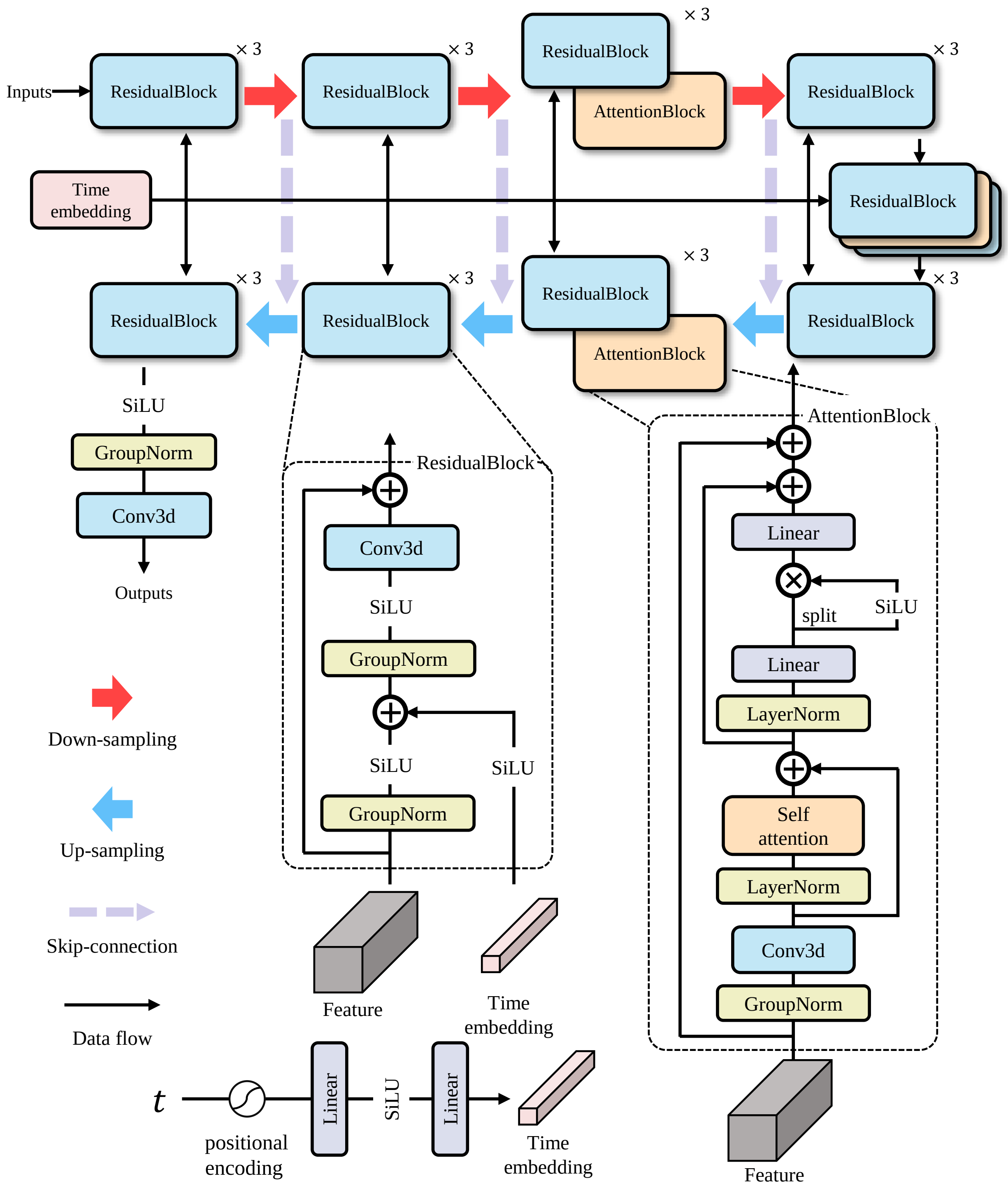


Figure S4: Details of the denoising UNet for CryoDiff.

CryoDiff is based on a conditional diffusion framework (Rombach *et al.*, 2022a,b), in which a time- and map-conditional 3D U-Net serves as the reverse denoising network. Specifically, we extend the time-conditional 2D U-Net backbone of latent diffusion models to volumetric cryo-EM density inputs by adopting 3D convolutions and incorporating map conditioning via channel-wise concatenation.

The network follows a hierarchical encoder–decoder architecture with four encoding stages, a central bottleneck, and four decoding stages, connected via skip connections. The base number of channels is set to 32 and increases progressively across scales. Residual convolutional blocks are employed at all stages to enhance local feature extraction. To capture long-range dependencies, attention modules (with 4 heads) are selectively introduced at the third encoder stage, the corresponding third decoder stage, and the bottleneck.

Temporal information is encoded using sinusoidal time embeddings, which are injected into the network through feature-wise modulation. This design enables effective integration of multi-scale structural information and conditioning signals during the denoising process. A detailed schematic of the network architecture is provided in Figure S4.

#### 3 Details of training data

We used a total of 980 non-redundant entries to train CryoDiff, which were randomly partitioned into training (737), validation (100), and testing (143) sets. The left panel of Figure S5 shows the resolution distribution of the CryoDiff dataset, while the right panel presents the distribution of entity types in the test set.

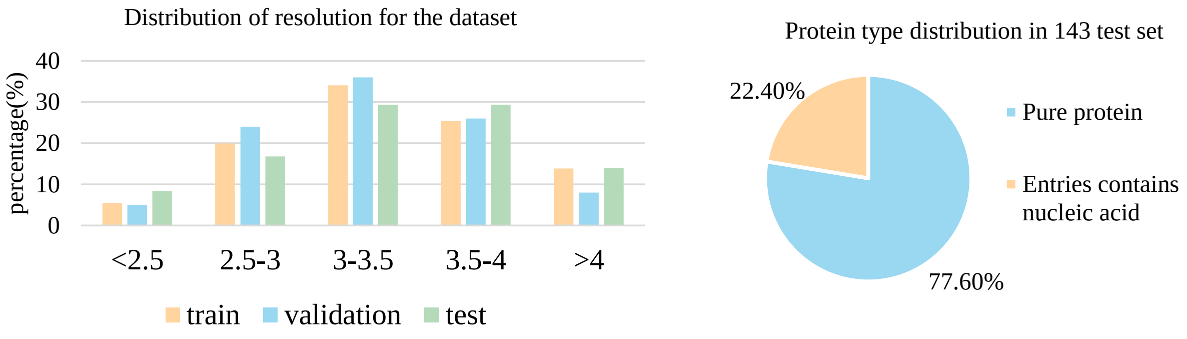


Figure S5: Distribution of resolution in the dataset and distribution of map composition in the test dataset.

### 4 Extended Evaluation of pMDD

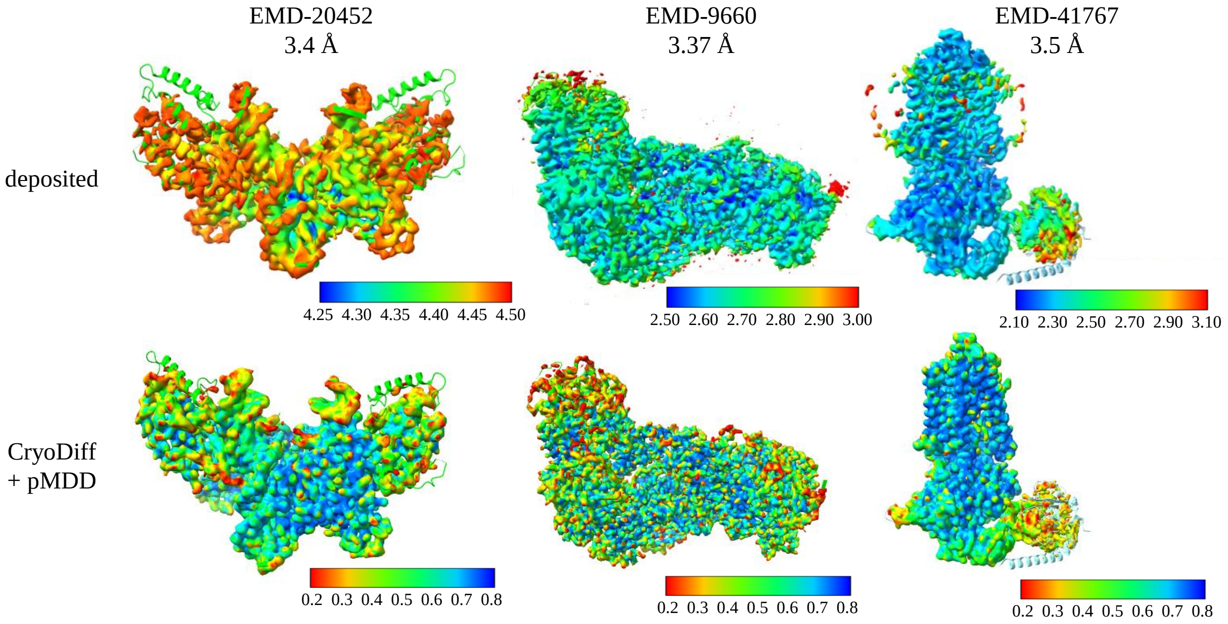


Figure S6: Cases shows pMDD results for EMD-20452 (3.4 Å), EMD-9660 (3.37 Å) and EMD-41767 (3.5 Å), Deposited maps colored with local resolution calculated from CryoRes are shown in the left section and CryoDiff-processed maps colored with pMDD values are shown in the lower section.

While the relationship between pMDD and Q-score demonstrates its ability to reflect model-based resolvability, Q-score itself relies on the availability of atomic models and therefore may not fully capture the intrinsic quality of the input maps. To further assess the general applicability of pMDD in a model-independent manner, we additionally examine its relationship with local resolution estimated directly from the deposited maps. As shown in Figure S6, we present three representative examples in which the deposited maps are colored by local resolution derived from CryoRes (Dai *et al.*, 2023) alongside the corresponding CryoDiff-processed maps colored by pMDD values. Across all cases, regions with poorer local resolution in the deposited maps consistently correspond to reduced pMDD values after enhancement. These results further support that pMDD provides a reliable indicator of local map quality.

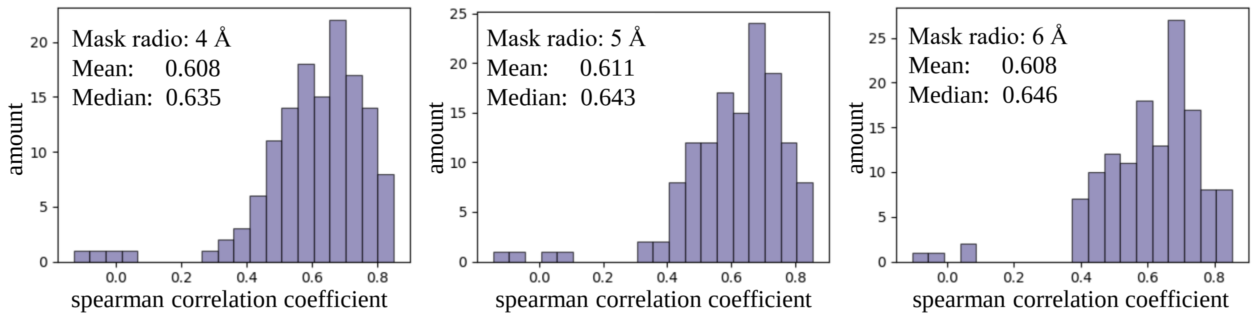


Figure S7: Distributions of spearman correlation coefficient between pMDD values and Q-score on 143 primary maps using different mask radios.

In addition, Figure S7 presents the distributions of Spearman correlation coefficients (SCC) between residue-wise pMDD values and Q-score across 143 test maps, evaluated using different mask radii. The results show a consistently correlation, with mean SCC values of 0.608, 0.611, and 0.608, respectively. These results further support pMDD as a reliable metric for reflecting the local resolvability of CryoDiff-processed maps.

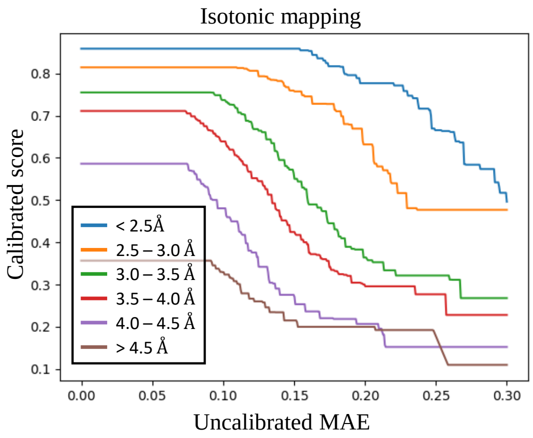


Figure S8: Monotonic mappings learned by isotonic regression across different global resolution bins.

To reduce the influence of global resolution on the relationship between raw uncertainty and local resolvability, we introduce a resolution-aware uncertainty calibration strategy. Specifically, voxel-wise raw MAE values are first computed from diffusion samples and paired with local Q-score as supervision signals. A total of 402 cryo-EM maps from the training dataset are used to learn this calibration. The dataset is stratified into multiple global resolution bins, and within each bin, isotonic regression is applied to learn a monotonic mapping between raw uncertainty and Q-score. The resulting set of bin-specific mappings is then used to transform raw MAE into a calibrated uncertainty measure (pMDD), ensuring consistency across resolution ranges while preserving the relative ordering of voxel-wise values. As shown in Figure S8, the learned isotonic functions exhibit clear resolution-dependent variations, indicating that the relationship between raw MAE and resolvability is not universal across different resolution regimes.

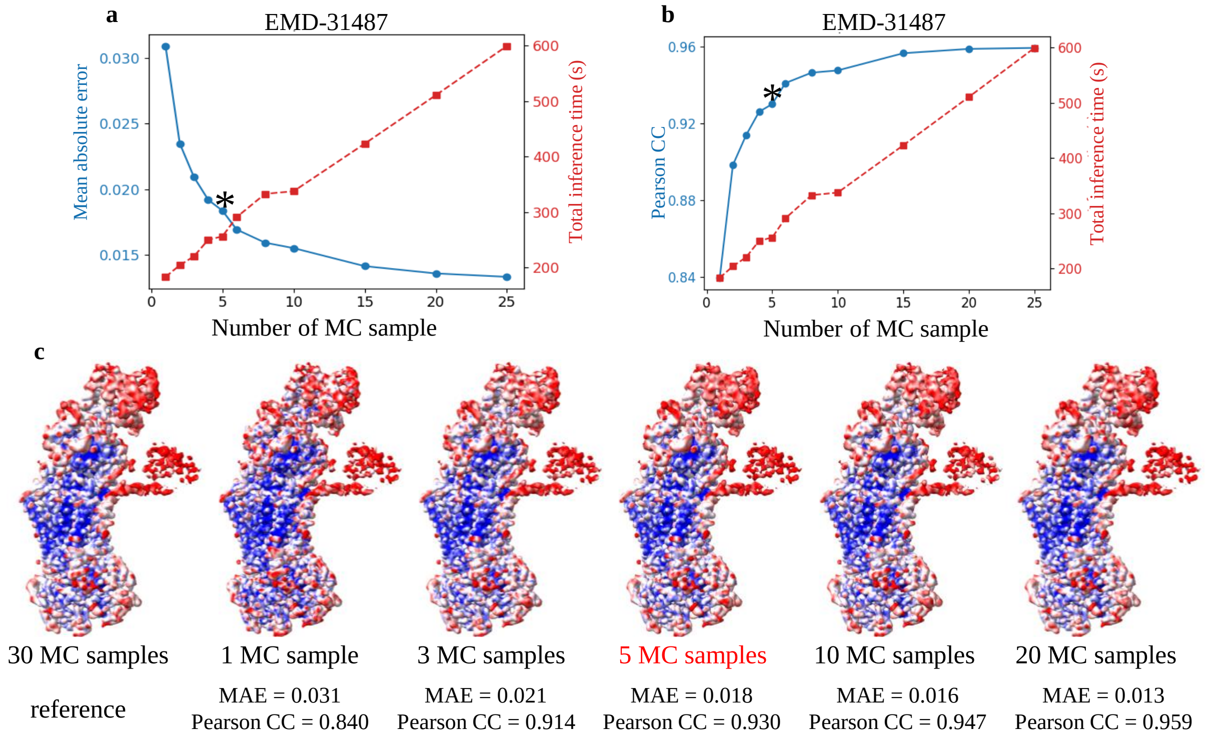


Figure S9: **Validation of the selection of the number of Monte Carlo (MC) samples. The chosen MC sample number is highlighted by an asterisk (*).** **a** Dual-axis plot showing the mean squared error (MSE) between pMDD maps obtained with different MC sample numbers and the reference pMDD map computed with 30 MC samples (left axis, blue dots), together with the total inference time as a function of the MC sample number (right axis, red squares). **b** Similar to a, but using the Pearson correlation coefficient (PCC) instead of MSE to quantify the consistency between pMDD maps. **c** Visual comparison of pMDD maps generated with different MC sample numbers, with the corresponding MSE and PCC values shown at the bottom.

To determine an appropriate number of Monte Carlo (MC) samples for reliable uncertainty estimation while maintaining computational efficiency, we performed a systematic validation on EMD-31487 (3.81 Å, PDB:7F7F) (Xu *et al.*, 2022). Specifically, pMDD maps generated with different numbers of MC samples were compared with a reference pMDD map computed using 30 MC samples, which was treated as an approximate converged estimate. The discrepancy between pMDD maps was quantified using the mean squared error (MSE) and the Pearson correlation coefficient (PCC), while the corresponding total inference time was also recorded.

As shown in Figure S8a and b, both MSE and PCC rapidly converge as the number of MC samples increases.

Meanwhile, the inference time grows approximately linearly with the number of MC samples. Visual inspection of pMDD maps (Figure S8c) further confirms that the qualitative patterns remain largely unchanged across different MC sample numbers. Based on these observations, we selected an MC sample number of 5 as a practical compromise between estimation accuracy and computational cost, which provides sufficiently stable pMDD estimates while significantly reducing inference time.
